# Scanning Single Molecule Localization Microscopy (scanSMLM) for super-resolution optical volume imaging

**DOI:** 10.1101/2022.04.01.486682

**Authors:** Jigmi Basumatary, Neptune Baro, Prakash Joshi, Partha Pratim Mondal

## Abstract

Over the last decade, single molecule localization microscopy (SMLM) has developed into a set of powerful techniques that has improved spatial resolution over diffraction-limited microscopy and demonstrated the ability to resolve biological features at the very molecular scale. We introduce a single molecule based scanning SMLM (scanSMLM) system that enables rapid volume imaging. Using a standard widefield illumination, the system employs a scanning based detection 4f-sub-system suited for volume interrogation. The 4f system comprises of a combination of electrically-tunable lens and high NA detection objective lens. By rapidly changing the aperture (or equivalently the focus) of electrically-tunable lens (ETL) in a 4f detection system, the selectivity of axial (*Z*) plane can be achieved in the object plane, for which the corresponding image forms in the image/detector plane. So, in-principle one can scan the object volume by just changing the aperture of ETL. To carry out volume imaging, a cyclic scanning scheme is developed and compared with conventional scanning routinely used in SMLM. The scanning scheme serves the purpose of distributing photobleaching evenly by ensuring uniform dwell time on each frame for collecting data (single molecule events) throughout the specimen volume. With minimal change in the system hardware (requiring an addition of ETL lens and related hardware for step-voltage generation) in the existing SMLM system, volume scanning (along *z*-axis) can be achieved. To demonstrate, we imaged fluorescent beads embedded in a gel-matrix 3D block as a test sample. Subsequently, scanSMLM is employed to understand clustering of HA single molecules in a transfected cell (Influenza A disease model). The system for the first time enables visualization of HA distribution in a 3D cells that reveal its clustering across the cell volume. Critical biophysical parameters related to HA clusters (density, #*HA/cluster* and clustered fraction) are also determined.

---

To be able to visualize distributed single molecule in a cell volume is an incredible feat. This gives a plethora of information related to local physiological in a cell and helps in deciphering the underlying biological mechanism. We demonstrate a scanning single molecule localization microscopy (scan-SMLM) that enables such a capability. The technique uses an electrically-tunable lens in the detection sub-system to locate single molecules in different layers of a 3D cell. The cells were transfected with photoactivable Dendra2-HA plasmid DNA that imitate infection (entry, replication and exit of viral particles) in an Influenza A model.

In the last decade, SMLM techniques have taken a big leap with resolution approaching sub-10 nm regime and temporal resolution of few milliseconds. Recent advances in SMLM such as, STED, MINFLUX, dSTORM, SIMPLE and ROSE have shown sub-10 nm resolution [2] [3] [4] [5] [6] [8]. With the advent of molecules that have high quantum yield and large molar extinction coefficient (e.g., pamCherry, Dendra2, Cy5, and Alexa Fluor 647), the resolution has bettered by an order. Non-the-less the fact that super-resolution is restricted to single plane of interest has somewhat handicapped the progress of SMLM. However, few techniques have shown promise for simultaneous multi-plane imaging, a step towards volume imaging. In this direction, the integration of light sheet and super-resolution has great potential. Specifically, 3D superresolution microscopy is reported by pairing tilted light sheet with long-range PSF [11]. The technique allows determination of the location of individual molecules detected throughout the volume excited by the light sheet. Moreover, the technique allows 3D SPT and single molecule SR imaging in 3D [10]. Off-late, localization microscopy over large volumes (upto 20 *µm*) is achieved by integrating lattice light sheet with cell-permeable chemical probes [11]. Moreover the technique also demonstrate super-resolution correlative imaging with photoactivable probes. Another promising technique is 3D STORM that uses optical astigmatism to determine both lateral and axial positions of individual molecules [12]. Multicolor STORM imaging of large volume has enabled ultrathin sectioning of ganglion cells to understand nanoscale co-organization of AMPA receptors and its spatial correlation with neuroligin-1 [13] [14] [15]. A recent technique primarily based on RESOLFT microscopy have shown 3D imaging of single molecules organization in a cell [16]. The technique has reported volumetric imaging of exosomes labeled with CD63-rsEGFP2 in live U2OS cells. Other techniques predominately based on evanescent light such as SMILE has demonstrated volume imaging capabilities in a single cell [17]. Although these techniques are promising, but they are not suitable for 3D imaging since they are primarily designed for 2D imaging. Moreover these techniques are complex and comes with a lot of constraints.

Classical SMLM techniques involves extracting single molecule signatures from a large set of recorded frames collected at video-rate. Subsequently, the bright spots are conveniently modelled by a 2D Gaussian, from which the centroid and number of photons are determined. These parameters serves as the basis to determine its position and size (governed by localization precision, 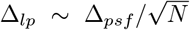where, Δ _*psf*_ and *N* are the diffraction-limited PSF and number of detected photons, respectively [18]). Since the invention of single molecule super-resolution technique (such as STED, fPALM/PALM, STORM), the technique has enabled many studies which were earlier thought to be nearly impossible [19–26]. Thereafter, many important variants have emerged that have harnessed its potential in a wide range of research disciplines. Specifically, the last decade has seen many variants including, ground-state depletion microscopy (GSDIM) [27], super-resolution optical fluctuation imaging (SOFI) [28], points accumulation for imaging in nanoscale topography (PAINT) [29] [30], simultaneous multiplane imaging-based localization encoded (SMILE) [31] [32], individual molecule localization–selective plane illumination microscopy (IML-SPIM) [33], MINFLUX [8],POSSIBLE microscopy [1] and others [34–41]. But seldom, techniques have demonstrated the ability to map 3D organization of single molecules in a cell. Therefore, methods that facilitate visualization of single molecules distribution in a cell volume are highly desirable. Such techniques expand the reach of super-resolution technique beyond traditional imaging and have the capability to take it forward.

In this article, we propose a scanning single molecule localization microscopy (scan-SMLM) that enables volume imaging. The technique employs a dedicated 4f detection system that comprises electrically focal-length tunable lens (ETL) which facilitate focusing of fluorescence originating from both focal and off-focal object planes on to the fixed image plane (detector plane). This is possible due to the ability of electrically-tunable lens to alter its focus by changing the shape of aperture, thereby facilitating both divergent and convergent rays (originating from off-focal planes) to focus on the camera (image/detector plane). The technique is demonstrated on bead sample, and further progressed to understand single molecule clustering in Influenza A disease model. In this respect, a patent for working scan-SMLM system along with associated optical design details is filed in India patent office [42].

We study the influenza type-A model that involves dynamic clustering of the glycoprotein Hemagglutinin (HA) in transfected NIH3T3 cells. HA protein is an antigenic glycoprotein found on the surface of influenza viruses and is responsible for binding the virus to the cell. Cells accumulate these proteins based on the local physiology during the onset of viral infection (Influenza A) [43] [44] [45]. It may be noted that HA clustering is a key process and have a direct bearing on the infectivity rate. So, it becomes essential to understand the HA accumulation process and find methods to disrupt it. In the present study, the cells were transfected with Dendra2HA plasmid DNA and cultured for 24 hours before fixing them for superresolution imaging. The photoactivable Dendra2 molecule involves the conversion between two states (on/off) mediated by triplet state. Accordingly, the priming or excitation of the anionic cis-chromophore populates the S1-state [46]. The de-population occurs via either fluorescence pathway or low-yield intersystem crossing to the lowest triplet state, T1. The mechanism involving conversion of primed state strongly relies on the creation of a triplet intermediate state. This results in fluorescence intermittencies such as triplet state transitions (popularly known as blinking) that are observed in many molecules. In general, several photoactivable fluorescent proteins show blinking timescale of a few tens of milliseconds (e.g., Alexa555 has a timescale of 22± 6 *ms* in deoxygenated PBS buffer [47], and Dendra2 has an average bleaching rate of 23.1*±* 1.9 *s*^-*1*^ [48]). This necessitates that the images be acquired between 35 - 40 frames / sec. While a significant fraction of photoswitches is non-fluorescent, a small subset is stochastically activated and localized. The molecules can be localized to high precision by fitting a Gaussian to their point spread function and determining the number of collected photons. SMLM techniques requires collection of several hundred frames containing single molecule signatures to reconstruct a single super-resolved 2D image. This leads to high precision localization of single molecules to render a super-resolution map of the specimen. Here, we propose cyclic scanning that facilitates even scanning all the specimen layers / planes in a periodic manner. This scanning configuration brings in uniformity across the planes and avoids long dwell time on a single plane as done in the conventional SMLM.

The schematic diagram of the proposed scanSMLM system is shown in Fig. 1. The illumination is achieved by a couple of lasers (405 nm light for activation and 561 nm light for excitation) and a high NA objective (Olympus, 100X objective, 1.3 NA). The resultant illumination PSF is marked by a black circle along with focal plane (*z* = 0) and off-focal planes (±Δ *z*). The emission from the focal plane (shown by blue rays) is collected by the objective lens and reflected by dichroic mirror to the a series of lenses that magnifies the image. Electrically-tunable lens (ETL) has a fixed focal length at a specific set current value that corresponds to image formation of z=0 plane. Now by increasing / decreasing current the aperture of ETL changes which focus conversing/diverging rays originating from off-focal planes to the detector / image plane. Aperture change is effected by changing current through the ETL which is controlled by an external voltage. So, the image formed at the detector plane is that of off-focal planes of the object. Hence, by just changing the external voltage, one can achieve z-scanning in the object plane. This is schematically shown in the inset of Fig. 1. Rapid scanning of z-planes in the object plane is achieved by a 4*i* driver and an automation circuit. The electrical lens driver 4i helps the user to control the focal power of ETL precisely. The lens current set into the firmware can be controlled by an analog applied voltage (0-5V) on the hardware Pin -B (see, Fig. 1). The applied voltage is linearly mapped in firmware to the range defined by the lower and upper current limits in the hardware configuration tab of Lens Driver Controller. The discrete step voltages are generated using digital potentiometer (MCP41010) via Arduino (microcontroller) (see, Fig. 1).

**FIG. 1:**
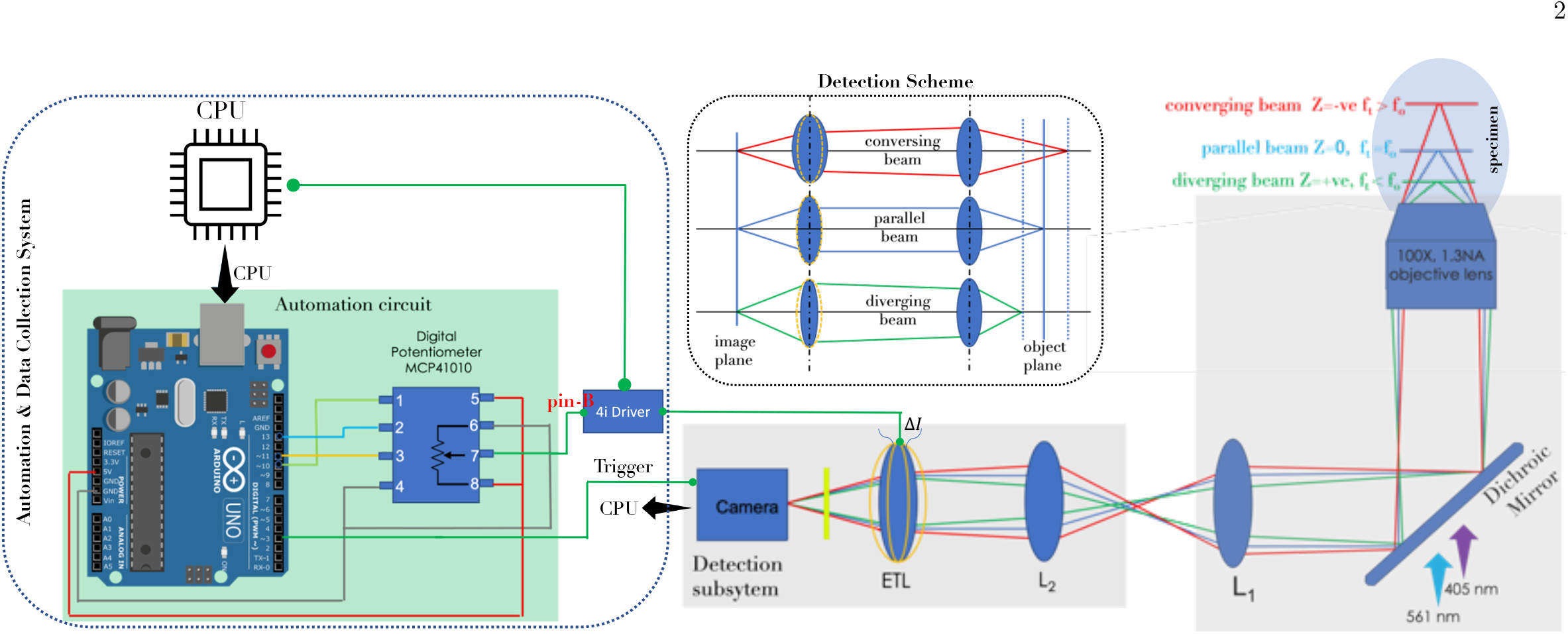
Schematic diagram for the proposed *scanSMLM* super-resolution microscopy. The illumination sub-system consists of activation laser (405nm) and excitation laser (561nm). The beams are combined by a dichroic mirror and focused to the specimen by a high NA objective lens (100X, 1.3 NA). The fluorescence from the single molecules are collected by the objective and directed to the EMCCD detector. On its way, the light is filtered by a set of filters. As additional magnification is introduced by a combination of biconvex and electrically-tunable lens. The electrical lens driver 4*i* is used to control the focal power of ETLs. Discrete voltage step generated using digital potentiometer is applied on pin B of 4i hardware driver which controls the lens current set into firmware. The process of voltage generation, ETL tuning and camera are synchronised via Ardino [9].

To facilitate bias-free scanning of all the object planes / layers, a cyclic scan technique is adopted as shown in Fig. 2A. The table displays #frame versus #planes with each element *P*_*ij*_ corresponding to the *i*^*th*^ plane undergoing *j*^*th*^ scan. In the cyclic scan scheme, all the planes are treated equally with all of them undergoing first round of interrogation (activation and excitation). This is represented by *P*_*i*1_ element in the scan-matrix. Similar process is carried out to acquire other scan elements column-wise i.e, *P*_*i*1_ followed by *P*_*i*2_ and so on till *P*_*iN*_, where *N* represents the total number of realization of a single plane. On the other hand, traditional SMLM (see, Fig. 2B) mostly follow conventional row-wise scanning of planes with first plane illuminated *N* number of times. This is represented by *P*_1*j*_. Sequentially, data for other planes are collected i.e, first plane elements *P*_1*j*_ followed by second plane elements *P*_2*j*_ till the last plane *P*_*Mj*_ is collected. Moreover, existing SMLM scanning biases against the planes and hence cannot account for uniformity. The corresponding digital potentiometer (DPM) output (that linearly maps the current range set in firmware) versus time diagram are also shown for scan-SMLM and conventional SMLM in Fig. 2(C,D). In addition, synchronized time line of EMCCD detector (exposure and readout time), DPM o/p and ETL tuning delays are also shown for volume acquisition cycles in Fig. 2E.. For comparison, conventional scan employed in traditional SMLM for collecting data plane-wise along with DPM voltage (for selecting specific plane) is also displayed (see, Fig. 2F).

**FIG. 2:**
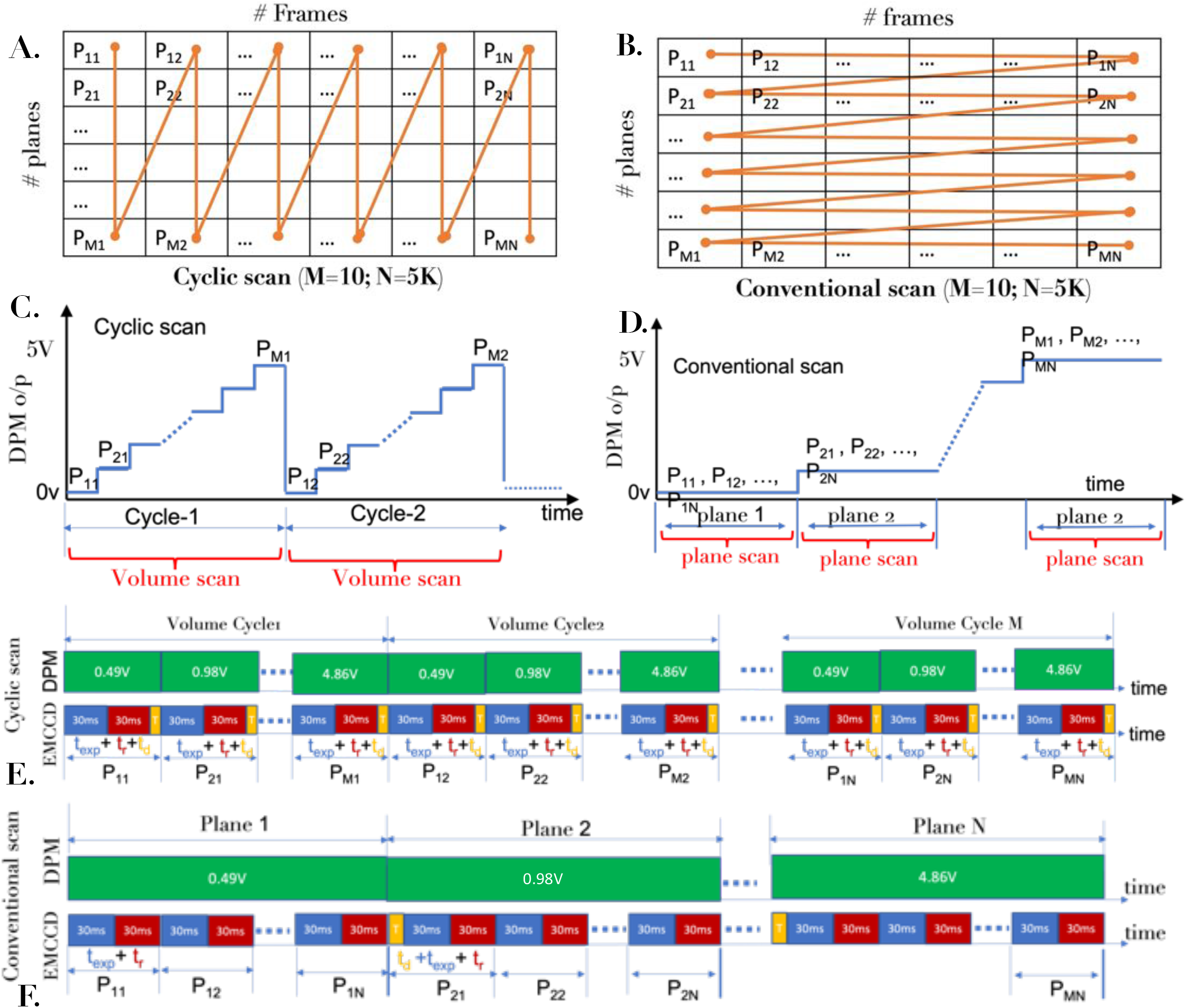
Volume scanning schemes: (A) Cyclic Scanning, (B) Conventional scanning. The element *p*_*ij*_ represents image of *i*^*th*^ plane during *j*^*th*^ cycle. (C) Generated step voltage for cyclic scan volume-wise, while (D) shows conventional scan plane-wise. In cyclic scanning mode for one step voltage, one frame of diffraction-limited single-molecule image is recorded. Whereas in conventional scanning mode for one step voltage required all the single molecules frames are collected for a particular plane before proceeding to the next plane. (E,F) The sequencing of exposure time, *t*_*exp*_; readout time, *t*_*r*_, and ETL response delay, *t*_*d*_ =5 *ms* for volume and plane scanning cycles employed in cyclic and conventional scanning, respectively. The time synchronization of EMCCD camera and DPM discreet output step voltage generation is also displayed.

For accurate data acquisition, the system needs to be calibrated across the illumination volume and repeatability need to be ascertained. Based on the *Z*-resolution (≈ 500 *nm*) of the inverted microscope, we have chosen to work with 10 planes. Subsequently, the specimen planes are calibrated with the focal length of the electrically-tunable lens as shown in Fig. 3B along with the enlarged inset that shows the region-of-interest (0-4.5 *µm*) beginning from 1100 nm from the coverslip. The characteristics of ETL lens at varying voltage or current is shown in Fig. 3C. Fig. 3A shows the schematic for calibrating scan-SMLM system using fluorescent nano-beads (size ∼170 *nm*, ex/em: 505/515, Invitrogen, USA). The beads were embedded in an agarose gel-matrix. Uniform distrubution of beads in the gel-matrix is ensured. A 3D block of the gel-matrix is cut and imaged using scan-SMLM (cyclic scan). The images were recorded and the corresponding raw data is shown in **Supplementary Video 1**. For reliable periodic scanning of specimen layers, we have correlated each plane with itself after a complete periodic scan i.e, between plane *i* and *i* + 10. The corresponding correlation plot is shown in Fig. 3D along with the **Supplementary Video 1**. A small variation in correlation ascertains the reliability of the system for cyclic or periodic scanning scheme. In addition, we have carried out drift measurement over the data collection time as shown in Fig. 3E. We noted a z-drift of ∼0.2 *nm/min* for the reported data. Along with other chosen parameters (z-sampling, region of interest and the distance from coverslip), the system is employed for recording single molecule data.

**FIG. 3:**
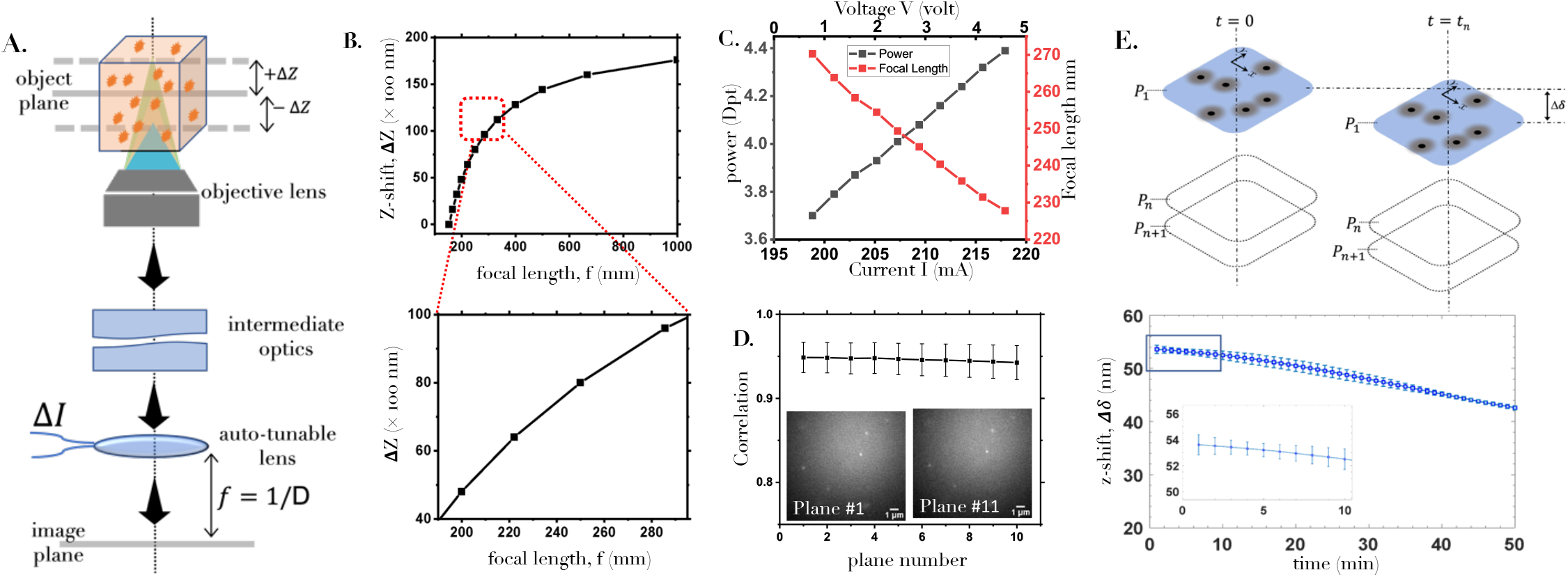
(A) Schematic of detection sub-system showing the connection between shift in object plane (Δ*Z*) and its image formation by the electrically-tunable lens in the detector plane (image plane). (B) The characteristics of z-shift (Δ*Z*) versus focal length (*f*) along with its operating region, 0 −4.5 *µm* in the specimen (see, enlarged section). (C) The corresponding plot relating power / focal length (with an offset of 150 *mm*), and current / voltage. (D) The correlation between *P*_*i,j*_ and *P*_*i*+10,*j*_ image of fluorescent beads of size 170‘*nm* (embedded in agarose gel-matrix) in a cyclic scan. The correlation between them is found to be *>* 94% demonstrating high repeatability of the planes. See **Supplementary video 1** that shows the accuracy of cyclic scanning method. (E) Axial drift measurement over data collection time.

Next, the scan-SMLM system is employed to understand and quantify HA clustering in a Dendra2-HA transfected (Influenza A model). The transfected cells were identified using a blue light (470-490 nm) and the data is recorded by a sensitive EMCCD camera. The ETL is synchronized with the camera which is operating at 36 *frames/s*. Each cycle consists of 10 images collected from respective Z-planes (plane 1-10) of a single cell with an inter-plane spacing of 500 *nm* as shown in Fig. 4A. Many such cycles are taken to obtain enough number of single molecule signatures throughout the volume (see, Fig. 4B). This sums up to a total of *M* × *N* images. For our study, we have taken *N* = 5000 cycles, and *M* = 10 planes. Technically, single molecule events can be quantified (its position and size) directly in a 3D volume, and the distribution of HA molecules can be obtained in a 3D cell volume. We have employed both conventional and cyclic scan for volume reconstruction. Few selected 2D reconstructed planes (#1, #5, #10) within the cell volume are shown in Fig. 4C. The corresponding average localization precision for both the scanning schemes are shown in Fig. 4D. In addition, photobleaching characteristics for both cyclic and conventional scan are displayed in Fig.4(E, G). It is evident that cyclic scan show a steady decrease in photobleaching whereas it varies for conventional scan depending upon the z-plane. The corresponding SNR follows a similar trend as shown in adjoining plot (Fig.4F). This is predominantly due to exposure of a single planes when data is acquired for a specific plane. Overall, the advantage of cyclic scan over conventional scan is evident in terms of signal-to-background ratio (SBR) and overall photobleaching.

**FIG. 4:**
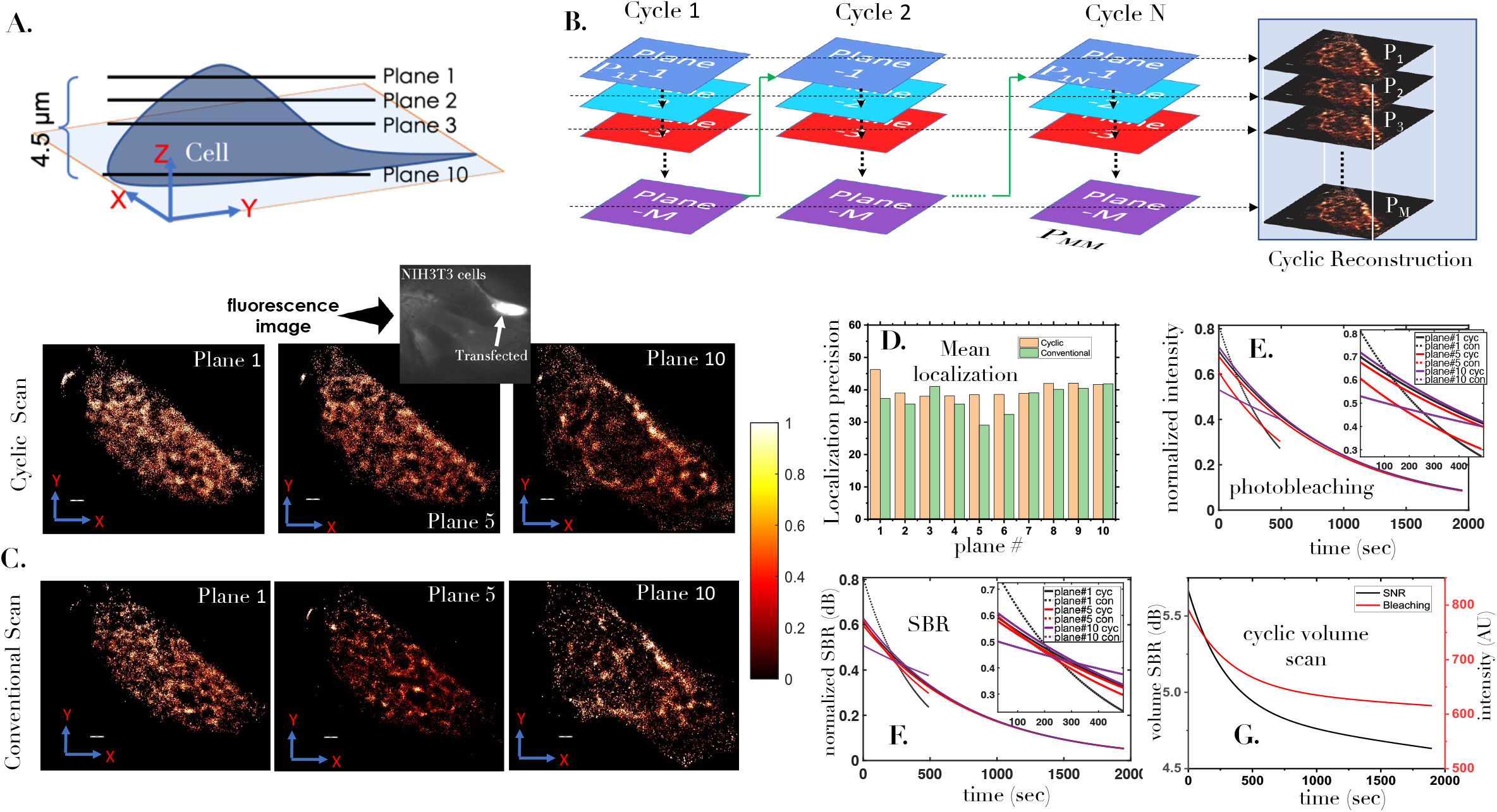
(A) A cartoon of 3D cell along with real fluorescence image of the transfected NIH3T3 cell. (B) Cyclic scanning for entire cell volume along with reconstructed images (*P*_1_ to *P*_*M*_). (C) Some of the sample super-resolution images (plane 1, plane 5 and plane 8) reconstructed from data acquited using cyclic and conventional scanning scheme. (D) The corresponding mean localization precision of all the reconstructed images (plane 1-10). (E) Plane-wise photobleaching study showing the decrease of fluorescence with time for both cyclic and conventional scan. (F) Signal-to-noise ratio (SNR) for cyclic and conventional scan. (G) Change in SNR and intensity with time for a cell volume. Scale = 1*µm*.

To determine critical biophysical parameters, we have employed both conventional SMLM and cyclic scanning scheme for which the reconstructed images are as shown in Fig. 5. The images are then subjected to cluster analysis. DBSCAN algorithm is used to identify the clusters and determine the parameters. Post 24 hours of transfection, the HA molecules are found to form clusters. We identified approximately, 45 clusters in a single plane. To better visualize the organization and close packing of HA molecules, we have shown reconstructed volume map in Fig. 6. It appears that the clusters are well connected across the layers of 3D cell. This is better visualized in the diagonal view and the related enlarged sections. In addition orthogonal sectional views (XY, XZ, YZ) are also shown that indicate direction-specific accumulation of HA molecules in the cell volume. Note that, the algorithm removes unclustered molecules that facilitates study of only the clustered molecules. A majority of the cluster size range from 0.18 −0.22 *µm*^2^ with an average cluster size of 0.2 *µm*^2^. The cluster density is known to be a critical indicator for accessing the progression of viral infection. The average density is found to lie between 950 −1050 #*M/µm*^2^. Surprisingly, the number of HA molecules per cluster varied from 160 - 200. These critical parameters are tabulated in the Fig. 6B for few chosen planes (*P* 1, *P* 5, *P* 10) of the transfected cell. Analysis on the entire cell volume indicate that a clustered molecule fraction (ratio of clustered to total HA molecules in the cell volume) of ∼43%. Another useful indicator is the cluster area fraction (ratio of the area occupied by the clustered molecules to the total area in a plane) which is found to be 9.85% per plane (average value). In addition, we have also observed a significant variation in the number of molecules per cluster per plane, while it is approximately 200 molecules per volume. The above parameters gives critical information related to rate of infection in a cell volume [49]. We anticipate that cell volume (considering all the planes) based biophysical parameter estimation facilitated by scan-SMLM may give a better understanding of HA clustering during Influenza-A viral infection.

**FIG. 5:**
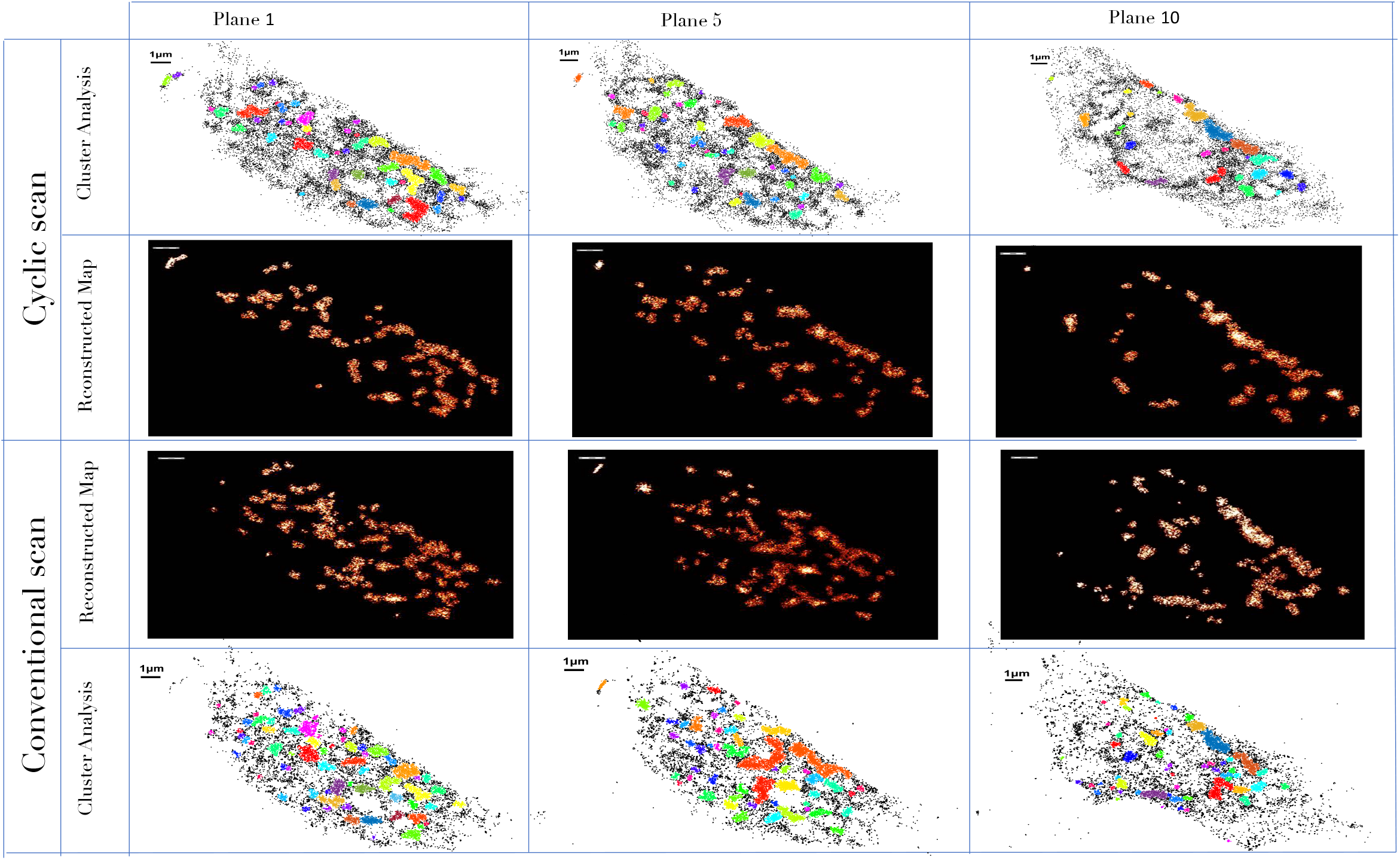
Reconstructed map of the distributed HA molecules in a transfected NIH3T3 cell for both cyclic and conventional scanning. DBSCAN algorithm is used to analysis clusters with parameters radius *‘* = 114 *nm* and minimum molecules of 35 for it to be designated as a cluster.Further analysis showing aggregated HA molecules in the cell.

**FIG. 6:**
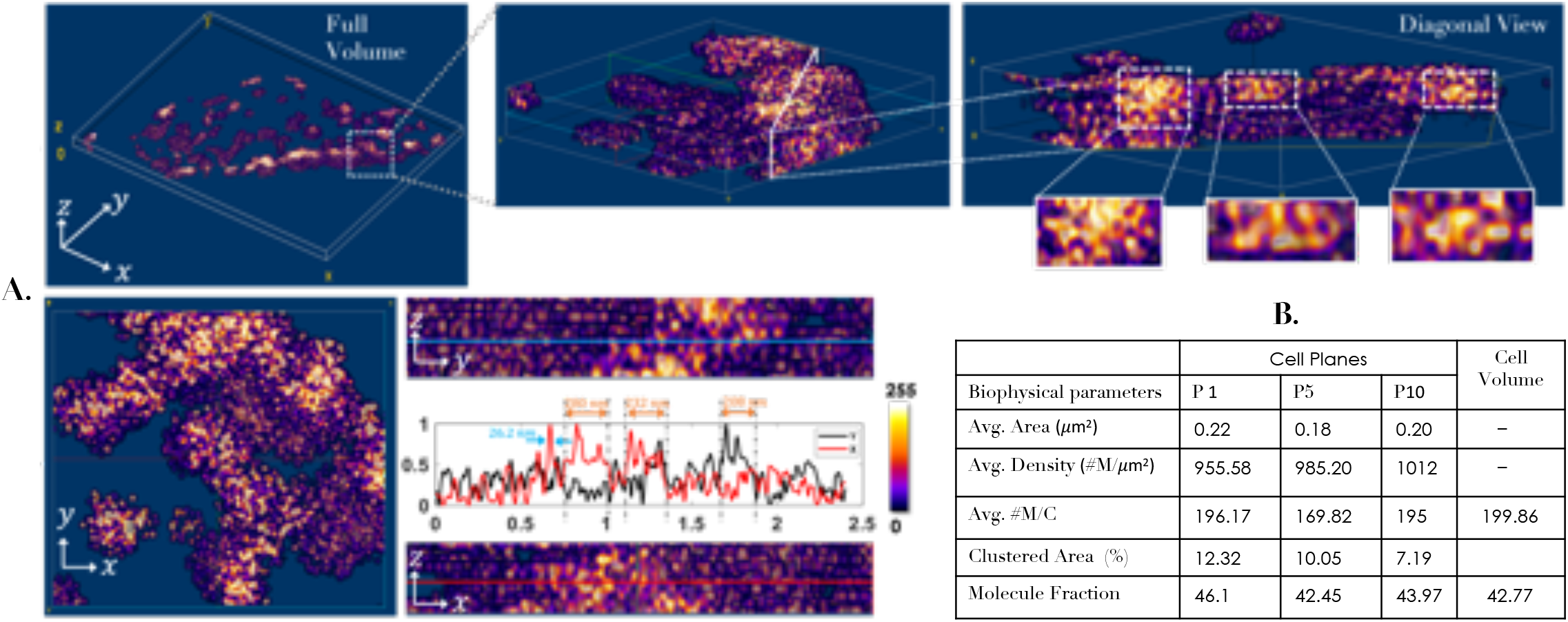
(A) Volume image of a cell displaying 3D organization of HA molecules. In addition, diagonal sectional view of the 3D cluster indicates the local density of clusters across the cell volume. Along side enlarged section of a single cluster is also shown. Orthogonal sectional views (along XY, YZ, XZ) of a specific region of 3D cell. Alongside intensity plots are also shown that indicate average HA cluster size of ≈240 *nm* (see, orange arrow) and typical single molecule size (localization precision) of 26.2 *nm* (see, blue arrow). (B) Estimated biophysical parameters (average cluster area, density and # molecules per cluster) related to local cell physiology indicates that most of the clusters are spread over less than an area of 200 *nm* with an average density of about 1*k* #*M/µm*^2^. These parameters are critical for understanding HA dynamics and help determine underlying biophysical processes. In addition, estimated from the cell volume shows the clustered molecule fraction of ∼42.77, indicating 42.77% of the total HA molecules are clustered, and rest remains unclustered, post 24 hrs of transfection. The clustered area fraction (which is the ratio of clustered area to the total cell area) is found be about 0.981%, indicating a *<* 1% area is occupied by clusters.

## Conclusion and Discussion

An optical scanning supersolution volume imaging system (scan-SMLM) is developed for determining single molecule distribution in a cell volume. Unlike existing SMLM technique that are capable of reconstructing a single plane, the proposed scan-SMLM system reconstructs the entire volume. The newly developed cyclic scanning technique periodically scans and records single molecules signatures from all the planes in an unbiased manner, and directly reconstructs the super-resolution volume map. Correlation study show high repeatability (> 94%) of the planes for long imaging cycles. In practice, this is achieved by using electrically-tunable lens for focussing fluorescence from respective z-planes (object) on to the detector, thereby enabling direct volume reconstruction.

To demonstrated the benefits of scan-SMLM technique, we employed it to understand HA protein aggregation in a Dendra2-HA transfected NIH3T3 cells (Influenza A model). The transfected cells were cultured for 24 hrs, fixed and then imaged by scan-SMLM system. Cluster analysis is carried out using DBSCAN algorithm, and critical biophysical parameters are quantified. Analysis show cluster characteristics in an entire cell volume rather than a single plane. The estimated parameter include average area, average density and average number of molecules per cluster in a plane which are estimated to be 0.20 *µm*^3^, 984.26#*M/µm*^2^ and 187#*M/cluster*, respectively. In addition, clustered molecule fraction (estimated from cell volume) show that approximately 43% of molecules are clustered post 24 hrs of transfection These biophysical parameters are critical since this indicate the local physiological activity of HA molecules leading to clustering process in a transfected cell, and has direct consequence on the rate of infection. We anticipate that scan-SMLM can be used to investigate the entire cell volume and determine volume-based critical parameters for other disease models as well.

## Acknowledgements

Financial funding from parent institute (Indian Institute of Science) is highly acknowledged. PPM and JB conceived the idea. JB, PPM, NB, PJ carried out the experiment. PPM prepared the sample. PPM wrote the paper by taking inputs from all the authors.

## Data Availability

The data that support the findings of this study are available from the corresponding author upon request.

## Disclosures

The authors declare no conflicts of interest.

## Notes

### Competing Interest Statement

The authors have declared no competing interest.

### Summary of Updates

1. Correction to small mistakes. 2. Correct the calibration scale.

